# A multilevel statistical toolkit to study animal social networks: Animal Network Toolkit (ANT) R package

**DOI:** 10.1101/347005

**Authors:** Sosa Sebastian, Puga-Gonzalez Ivan, Hu Feng He, Zhang Peng, Xiaohua Xie, Sueur Cédric

## Abstract

How animals interact and develop social relationships regarding, individual attributes, sociodemographic and ecological pressures is of great interest. New methodologies, in particular Social Network Analysis, allow us to elucidate these types of questions. However, the different methodologies developed to that end and the speed at which they emerge make their use difficult. Moreover, the lack of communication between the different software developed to provide an answer to the same/different research questions is a source of confusion. The R package Animal Network Toolkit (ANT) was developed with the aim of implementing in one package the many different social network analysis techniques currently used in the study of animal social networks. Hence, ANT is a toolkit for animal research allowing among other things to: 1) measure global, dyadic and nodal networks metrics; 2) perform data randomization: pre-network and network (node and link) permutations; 3) perform statistical permutation tests. The package is partially coded in C++ for an optimal coding speed, and it gives researchers a workflow from raw data to the achievement of statistical analyses, allowing for a multilevel approach: from individual position and role within the network, to the identification of interaction patterns, and the analysis of the overall network properties.

## INTRODUCTION

Social Network Analysis (SNA) has caught much attention over the last decades in animal research. This graph theory discipline is a statistical framework that allows the study of interconnected systems. In biology, such systems are found at all organization levels, from molecules to ecosystems [1-9] making thus SNA techniques powerful multidisciplinary tools. SNA are not only multidisciplinary but also allow to study the systems at different levels (from node position to overall structure). The amount of SNA tools is so vast (one mode network [10], multiplex network [10], ERGM [11, 12], ego-networks [13], network metrics [14], network medialization [15], Network Based Diffusion Approach [4], etc.) that it is impossible to draw up an exhaustive list and to gather all of them into one sole software. This tremendous diversity of analytical techniques raises the issue of the pertinence and utility of some of the SNA tools used in the analysis of animal social systems.

Moreover, statistical analysis techniques have been developed in animal research to handle intrinsic biases when collecting data from animals (such as heterogeneity in individuals observations, possible misidentification, differences in groups size or group composition) and depending on the protocol for data collection (focal sampling, scan sampling, group fellow) [16]. In animal research, protocols are now well established through Null Models approach (NM) based on different types of randomization techniques [17]. The necessity to adapt the Null Models to the type of data collected is another limitation in the different SNA software currently available.

The motivation behind Animal Network Toolkit (ANT) comes from these limitations. We developed this R package in order to provide researchers in animal social networks (usually small networks under 1000 nodes) that have specific observational data protocols with a unique software to process their raw data and help them with the selection of the most appropriate metrics and statistical analyses. This paper outlines the large spectrum of analyses allowed by ANT:

1. A multilevel approach through the study of nodes, dyadic and global network metrics.
2. Appropriate randomization techniques (pre- or network permutations) depending on the data.
3. Large amount of networks metrics available, some commonly used in animal research and others lesser known.
4. Several statistical tests (correlation test, t-test, General Linear Model, General Linear Mixed Model) with the calculation of permuted p-values.
5. The possibility to analyze static or temporal networks.

ANT is currently available in open beta test version on the following GitHub web page: https://github.com/SebastianSosa/ant

## PACKAGE OVERVIEW

### ANT analytical protocols

ANT allows two different analytical protocols: 1) single network analysis, 2) multiple networks analysis. Single network analysis is the common analytical approach found in several studies where only one network (one group at one moment) is analyzed [18, 19]. Multiple networks analysis allows the user to examine similarities or differences between individual node metrics (see section *Network metrics*) between different groups and/or time-repeated measures on a single/multiple group(s) [20, 21]. ANT analytical protocols are synthesized in Figure I.

**Figure I.**
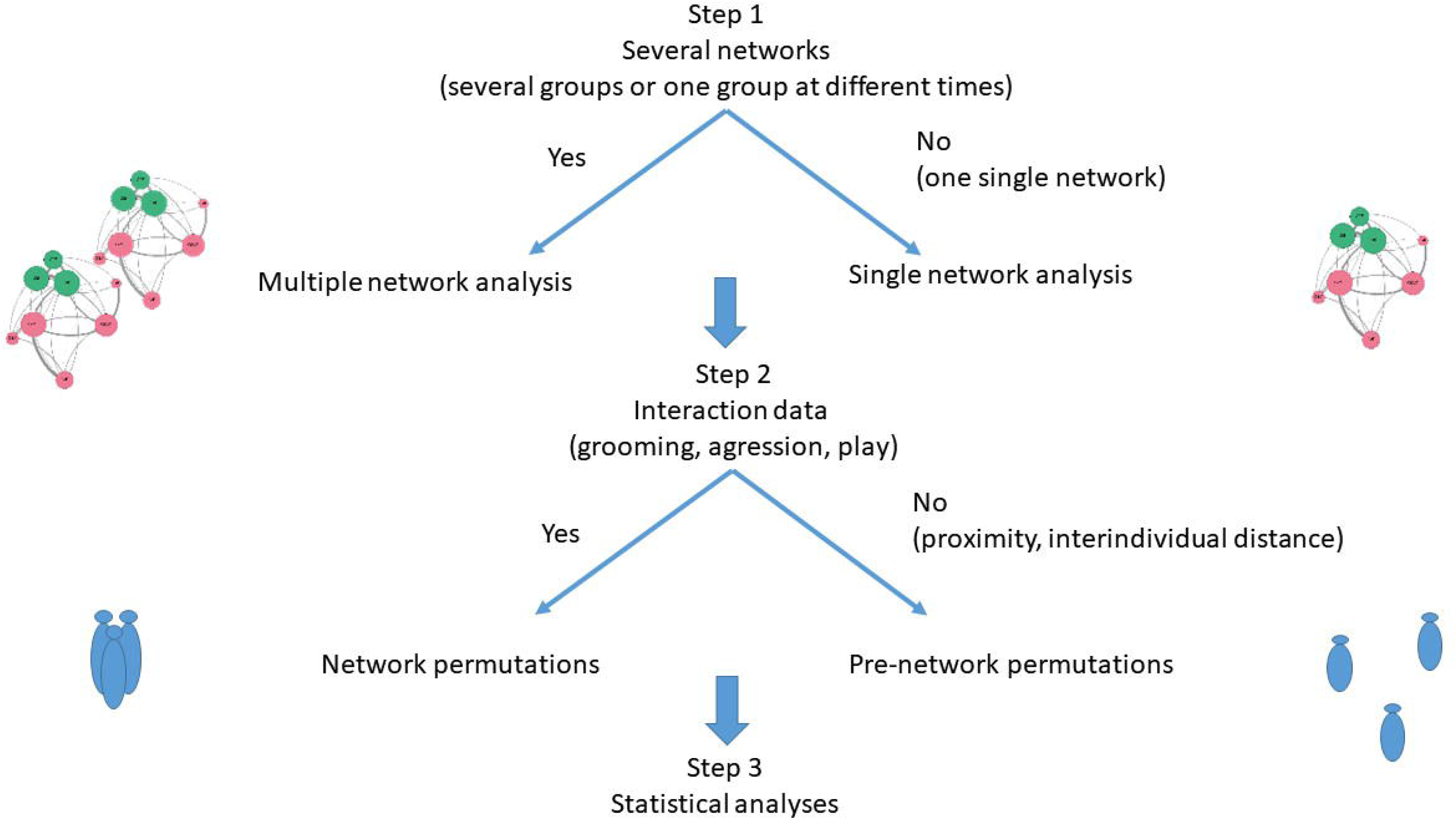

### Data input

ANT can process two types of data: 1) interactions (grooming, aggression, play, etc.) or associations data (interindividual distances), 2) data based on individual attributes (sex, age, dominance rank, etc.). This makes it possible to permute and/or to compute network metrics on interactional data and to store them in tables with individual characteristics to run permutations or statistical tests depending on the approach chosen for the NM.

Data based on individual attributes are generally data frames with each row representing an individual and each column representing a specific attribute such as sex, age or dominance rank etc. Combined with the individuals’ node metrics, it is then possible to study an individual’s centrality, the number of alters (congeners with whom the individual interacts) or its activity according to the individual attributes, and/or to extract patterns common to individuals with similar attributes.

Interactions/associations data can be represented in two different ways, as an adjacency matrix or as a table. An adjacency matrix is a square matrix ***X*** with *m* rows and columns,: ***X****=m*m*. Each column and row represent a node, with column 1 and row 1 corresponding to the same node. A directed network is represented by an asymmetric square matrix with the focal and emitting node (giver) in the row and the receiver in the column. An undirected network is represented by a symmetric square matrix in which the upper triangle (the triangle above the matrix diagonal) and the lower triangle (triangle below the matrix diagonal) are identical. The weight of the links is represented by numerical values in the matrix cells (in the case of a binary network, these cells consist of zeros, when the links is absent, and ones, when the link is present). In an interaction table, each row generally represents associations or interactions between two or more individuals, additional columns can indicate factors such as the date, the day, or the geographic location etc. In ANT, interaction tables can be converted into matrices to compute network metrics, or they can be used to realize pre-network permutation approaches (for more details on pre-network permutation approaches, see section Permutations).

When realizing multiple networks analytical protocol, the user has to import the different interaction/association data representing the different networks in an R format list, and optionally, the different data based on individual attributes for each of the network in an R format list to.

### Permutations

When examining inter-individual interactions within a group or a population, it is assumed that neither all the interactions nor all individuals are observed, that the times of observation vary from one individual to another, and that the data collected are intrinsically dependent. For these reasons, it is necessary to run permutation tests, as not all inferential statistical tests can be trusted. The Null Model approach (NM) is one of the many existing possibilities for the study of social networks.

The NM approach consists in creating random data sets from the observed data. The metric of interest *X* of the observed data and the posterior distribution obtained from the random data sets are then compared to examine if the observed metric X is significantly different from the random distribution by assessing the proportion of random values that differ from the observed values [16, 17, 22]. NM approach can be categorized according to the elements of the network that are being resampled (links or node labels), and according to whether they are built on raw data (before building the network: prenetwork randomization) or on the network (after building the network: network randomization) [22]. The permutation procedure has to be chosen according to the type of behavior collected and the protocol used for data collection. All of the permutation approaches available in ANT are in the family function ‘*perm’* with two sub-classes ‘*perm.ds*’ for permutation through data stream (pre-network) and ‘*perm.net*’ for permutations through network approach.

Pre-network randomization consists in permuting the alters of ego (the individual being examined), generating a network in which individuals interact randomly [16]. The alters are permuted according to the data sampling protocol used. So randomization can be restricted to specific periods, such as day time periods, geographical locations, etc. The specific control factor used may depend on the research question and the protocol used to collect data. Pre-network randomization is mainly used in ecology when inter-individual links are inferred by individual associations (interindividual distances or proximities), i.e. the ‘gambit of the group’ [17]. This method relies on the likelihood that individuals who frequently gather will also interact with each other. It is then possible to infer preferential interactions from the presence/absence of certain individuals. For this method, it is important to consider not only the size of the group observed, but also the difficulty of observation of specific individuals, which finally influences their frequency of observation. Thus, pre-network randomization is generally used for association data with a high risk of randomness (inter-individual distances) [17]. ANT allows two types of pre-network permutations: ‘group following’ and ‘focal sampling’. Each of these requires a specific data frame format and can be run with or without control factors.

Network permutations are used when examining direct behavioral interactions (grooming, agonistic interactions, etc.) and when groups are well defined [17]. This approach consists in randomizing link and/or node labels in a network. Node labels permutations (*i.e.* permuting nodes’ characteristics between each other) preserve the structure of node links. The goal of this approach is to determine whether a node’s position is linked to its intrinsic attributes. These permutations can be run on a data frame that gathers node labels and node network metrics that is being studied. Link permutations (*i.e.* permutation of the links between the nodes) are mainly used for the analysis of interactional patterns, *i.e.* how interactions are arranged and which individuals are involved.

For more details on the different permutations and their applications depending on the protocol used for data collection and the type of behavioral data collected, see Bejder [23], Whitehead [24], Whitehead [25], Croft [26], Farine [16], Sosa [22].

### Network metrics

Three types of network metrics according to the level of organization can be identified: global metrics, dyadic metrics, and node metrics. In ANT, all these metrics are grouped under the functions ‘*met’*.

Global metrics allow to study the overall network and to obtain valuable information regarding network efficiency, resilience, clusterisation, etc. In animal research, it remains difficult to analyze this type of metrics because several of them are strongly dependent with a basic metric called density (network ratio of observed links to all potential links). Density is sensitive to the sampling effort which in turn is linked to group size, making then very difficult to compare global metrics of different networks with no specific analytical protocol established [27-29]. All the global metrics available in ANT are synthesized in Table I. However, at the present time, it does not allow to compare global metrics across networks. Nevertheless, it is possible to test the significance of these metrics by running permutation tests.

**Table I.** Available ANT metrics and their short description.

| Metric level | Metric | ant code | Definition |
| --- | --- | --- | --- |
| Node | Degree | met.degree | Number of links |
|  | Outdegree | met.outdegree | Number of outgoing links |
|  | Indegree | met.indegree | Number of incoming links |
|  | Strength | met.strength | Sum of the weights of links |
|  | Outstrength | met.outstrength | Sum of the weights of outgoing links |
|  | Instrength | met.instrength | Sum of the weights of incoming links |
|  | R-index | met.ri | Ratio of outgoing and incoming links |
|  | Disparity | met.disparity | Variation of the weights of the links of a node |
|  | Eigenvector | met.eigen | Node centrality considering the connections of a node and its alters |
|  | Reach | met.reach | Node centrality considering the overall strength of a node alters |
|  | Affinity | met.affinity | Node centrality considering the overall strength of a node and that of its alters |
|  | Laplacian centrality | met.lp | Importance of a node in network cohesion |
|  | Betweenness | met.betweenness | Number of times a node lies within the shortest paths* of all pairs of nodes in the network |
| Dyadic | Assortativity** | met.assortativity | Degree to which a node is linked to others with the same attribute rather than linked to others with a different attribute |
|  | Geodesic distance | met.geodesic | Shortest paths* between each pair of nodes in the network |
| Global | Density | met.density | The number of actual connections divided by all potential connections in the network |
|  | Diameter | met.diameter | Greatest shortest path* |
|  | Global efficiency | met.ge | Efficiency of information transmission within the network |
|  | Centralization index | met.ci | Degree of network centralization |

\* Shortest distance (based on the number of links or weight of the links) connecting a given pair of nodes
\*\* The calculation of this metric is based on dyadic links but the value return applies to the whole network

Dyadic metrics allow the study of interaction patterns between individuals. These metrics inform how individuals interact according to their attributes. At the present time, ANT only allows the calculation of assortativity. All the dydaic metrics available in ANT are synthesized in Table I

Node metrics are the most frequently used metrics in animal research. They inform, among other things, about the centrality of an individual, the number of alters it has, or its activity according to individual attributes, and they reveal patterns common to individuals with similar attributes. Among the great diversity of node metrics we have chosen metrics that are frequently used in animal SNA research and some of them may represent promising tools to reveal new aspects (e.g. individual role in network cohesion) in future studies, such as the Laplacian centrality [30] or the normalized betweenness [31]. All the node metrics available in ANT are synthesized in Table I.

For more details on the different types of metrics, their mathematical formula, their interpretation, and their past use in animal research, see Whitehead [2], Sueur [32], Sosa [22] and ANT documentation.

### Statistical tests and results

The different steps to follow in ANT when applying an analytical protocol with pre-network permutations are: 1) perform permutations, 2) calculate the metrics, and 3) run the statistical tests chosen on all permutations. When applying an analytical protocol with network permutations, the steps to follow are: 1) calculate the metrics, 2) perform permutations, and 3) run the statistical tests chosen on all permutations. All the statistical tests available in ANT are in the family function ‘stat’. The available tests are: correlation test ‘*stat.cor’*, t-test ‘*stat.t*’, Linear Model (LM) ‘*stat.lm*’, Generalized Linear Model (GLM) ‘*stat.glm*’, Generalized Linear Mixed Models (GLMMs) ‘*stat.glmm*’, assortativity test ‘*stat.assortativity*’, and TaurK correlation ‘*stat.Taurk*’. Depending on the test selected, ANT automatically opts for the metric that will be extracted for the observed and each permutated network in order to calculate the permuted p-value (correlation coefficient for correlation test, the t student for student test, and the estimate of the different factors for LM, GLM and GLMM).

The output is adapted to the type of test run and the output is different according to each one of them but one common element is always found: the posterior distribution of the statistic of interest. The function ‘ant’ allows to obtain the statistical results of permutation tests after the function of type ‘stat’. ‘ant’ function outputs are adapted to the permuted statistical test run. However, some outputs are common to all tests. The function returns a data frame with the statistics of the permutation tests:

1. P-values, by the nature of the test, they are one-tailed p-values, on the right or left of the distribution.
2. Measure of the ‘effect size’ of the posteriori distribution according to the statistics of interest: 95% confidence interval and the mean of the distribution a posteriori [33].
3. A histogram of the post-distribution of the statistics of interest obtained from the permutations and with the observed value highlighted in white.

Besides, for LM, GLM and GLMM, the function ‘ant’ returns a list of 3 elements:

1. The original LM/GLM/GLMM with the estimates, the standard errors, as well as the permuted p-values, the mean of the posterior distribution and the “95 confidence interval” of each factor.
2. The posterior distribution of the estimate of each factor
3. A vector of integers indicating the permutations that returned model errors or warnings (e.g. model convergence issues) and for which new permutations were done.

The use of a permutation approach may lead some models to encounter convergence and/or optimization issues when running specific permutations. This can also originate from the complexity of the model specified in regards to the data set. While running the first model (on the original data set), ANT checks for errors and warnings and asks the user if (s)he wants to continue. While computing the model on the permuted data, each time ANT finds a warning or an error, a new permutation is run until the warning and error message disappears. Two considerations are required at this point:

1. The function can be stuck in an infinite loop when the model always returns error and/or warning messages. Progress bar for progress visualization can help to detect such issue.
2. By repeating the permutation process on the data each time an error and/or a warning is indicated, the posterior distribution will be forcibly restrained within a specific range of values.

This is why the ‘*stat.lm*’, ‘*stat.lm*’ and ‘*stat.glmm*’ function will return the type of error or warning found while computing the models and the id of the permuted data set that caused the error and/or warning. The user has then four options:

1. Find the origin of the errors or warnings, create a new simple model, scale and/or center variables, check singularity, or many other possibilities as explained in Ben Bolker github web page: https://rstudio-pubsstatic.s3.amazonaws.com/33653_57fc7b8e5d484c909b615d8633c01d51.html
2. Compute diagnostic tests on the model with ‘ant’ function.
3. Use the option ‘*control = [g]lmerControl(calc.derivs = FALSE)*’ that turns off the derivative calculation performed after optimization. By doing so, most of the warnings can be discarded as explained in lme4 vignette:lme4 Performance Tips: https://cran.rproject.org/web/packages/lme4/vignettes/lmerperf.html

As such concerns have not been addressed in the literature, the choice of the most appropriate option is left to the user.

## DISCUSSION

ANT provides researchers with an all-in-one software in which they will find the analytical tools most commonly used in animal social network research. ANT is a user-friendly software that enables beginners to run complex analyses on social networks. Besides, ANT also provides users with a wide range of functions that opens up countless possibilities for those who have greater programming and analytical skills and want to make a more autonomous use of these tools. We hope that this will enable users to focus less in coding and more on selecting the most appropriate permutation approach, statistical test(s) and metric(s), and to reflect more deeply on why this metric (or metric version) rather than another. It is still frequent to see biased studies because the lack of questioning on the most appropriate analytical tools to use when performing an analysis. By giving access to global, dyadic and node metrics, our aim is to enable users to adopt a multilevel approach and thereby understand the position of individuals in a group, the patterns of interaction, and the impact of these two levels on the global network structure [22]. We hope that ANT will represent an analytical standard for upcoming studies on animal social networks. We will try to update the package as often as possible to add new metrics and analytical techniques. ANT is a long-term project that is meant to evolve and improve. In the future, it may give access to additional techniques used in other research fields such as population ecology, as well as additional permutation tests and metrics. As this is a multi-collaborative project, we encourage everyone to take part in the development of future versions of ANT.

